# Experimental copper exposure, but not heat stress, leads to elevated intraovarian thyroid hormone levels

**DOI:** 10.1101/717157

**Authors:** Ruuskanen Suvi, Mottola Giovanna, Anttila Katja

**Author notes:** Corresponding author: Suvi Ruuskanen, Department of Biology, University of Turku, Vesilinnantie 5, 20500 Turku, Finland.

## Abstract

Climate change and pollution are some of the greatest anthropogenic threats to wild animals. Transgenerational plasticity – when parental exposure to environmental stress leads to changes in offspring phenotype – has been recently highlighted as a potential mechanism to respond to various environmental and anthropogenic changes across taxa. Transgenerational effects may be mediated via multiple mechanisms, such as transfer of maternal hormones to eggs/fetus. However, sources of variation in hormone transfer are poorly understood in fish, and thus the first step is to characterize whether environmental challenges alter transfer of maternal hormones to eggs. To this end, we explored the genetic and environmental variation (in response to temperature and endocrine disrupting copper) in maternal thyroid hormone (TH), transfer to offspring in a common fish model species, the three-spined stickleback (*Gasterosteus aculeatus*) using multiple approaches: (i) We compared ovarian TH levels among six populations across a wide geographical range in the Baltic Sea, including two populations at high water temperature areas (discharge water areas of nuclear power plants) and we experimentally exposed fish to (ii) environmentally relevant heat stress and (iii) copper for 7 days. We found that populations did not differ in intraovarian TH levels, and short-term heat stress did not influence intraovarian TH levels. However, copper exposure increased both T4 and T3 levels in ovaries. The next step would be to evaluate if such alterations would lead to changes in offspring phenotype.

Capsule: We show that experimental copper exposure, but not heat stress (experimental or among-population variation), leads to elevated ovarian thyroid hormone levels in sticklebacks.

## Introduction

Climate change and pollution are some of the greatest anthropogenic threats to wild populations. Organisms may respond to changes in their environment by showing plastic responses (Habary et al., 2017; Parmesan, 2006; Stillman and Armstrong, 2015). One recently highlighted form of plasticity is transgenerational plasticity, i.e. when variation in parental environment leads to changes in offspring phenotype (e.g. Donelson et al., 2012; Donelson et al., 2018; Metzger and Schulte, 2017; Meylan et al., 2012; Salinas and Munch, 2012; Shama et al., 2014; Shama and Wegner, 2014). For example, Shama et al. (2014) showed in their seminal paper rapid transgenerational acclimation to elevated temperature, although the molecular mechanisms are not yet understood. Transgenerational effects may be mediated via multiple mechanisms, such as epigenetic markers or transfer of maternal hormones or RNAs to eggs/fetus (e.g. Adrian-Kalchhauser et al., 2018; Best et al., 2018; Kim et al., 2019; Metzger and Schulte, 2017; Meylan et al., 2012; Ruuskanen and Hsu, 2018).

Hormones transferred from the mother to embryos and eggs are known to profoundly influence offspring development, physiology, morphology, behavior and even survival across taxa (mammals, Dantzer et al., 2013; fish, McCormick, 1999; birds, Ruuskanen, 2015; Ruuskanen and Hsu, 2018; reptiles, Uller et al., 2007). One class of these hormones is thyroid hormones (THs, prohormone thyroxine T4, and biologically active tri-iodothyronine, T3) and recent studies suggest that maternal THs transferred to eggs and embryos are important for offspring development across vertebrates (fish, Brown et al., 2014; birds, Hsu et al., 2017; Hsu et al., 2019; mammals, Patel et al., 2011; Ruuskanen et al., 2016a). THs are a key class of hormones that control and regulate vital biological processes such as thermogenesis, reproduction but also growth and metamorphosis (Norris and Carr, 2013). THs are critical regulators of thermal acclimation in fish, increasing in higher temperatures, (e.g. Little et al., 2013) and plasma THs have been found to fluctuate with varying water temperature (e.g. Arjona et al., 2010; Cyr et al., 1998; Eales, 1985). However, THs are subject to endocrine disruption by various chemicals, such as PBC, dioxins and heavy metals, such as lead (Matthiessen et al., 2018; Norris and Carr, 2006). A less studied, but a potential endocrine-disrupting chemical (EDC) is copper (Cu), although the results are controversial: Copper exposure has been found both increase and decrease plasma THs in fish, depending on species and timing of exposure (Eyckmans et al., 2010; Hoseini et al., 2016; Oliveira et al., 2008).

Surprisingly, studies characterizing the environmental or genetic sources of among-female (or within-female) variation in egg TH levels are rare (Ruuskanen and Hsu 2018), yet, such variation could contribute to variation in offspring phenotype and fitness and help to understand the scope for transgenerational plasticity. Thus, the first step is to characterize whether environmental challenges affect transfer of maternal hormones to eggs. In birds, egg TH levels vary with food availability (Hsu et al., 2016) and temperature (Ruuskanen et al., 2016c), while T3 (but not T4) also shows heritable variation (Ruuskanen et al., 2016b, Hsu et al. 2019). In fishes, there is indication for variation in egg THs among stocks (rainbow trout, *Oncorhynchus mykiss,* Leatherland et al., 1989), which could reflect either genetic or environmental variation. Interestingly, McComb et al. (2005) reported that the egg T3 and T4 concentration in bonnethead sharks (*Sphyrna tiburo)* from Tampa Bay was consistently higher than from eggs from Florida Bay. The authors suggested that this may be due to higher temperatures in Tampa Bay, and speculated that egg THs might, thus, explain the faster growth rates and metabolic rates at this site. In the rare examples on endocrine disruption, experimental exposure to pollutants (lead, polybrominated diphenyl ethers and bisphenol A) resulted in decreases plasma and egg THs in zebrafish (*Danio rerio*) (Chen et al., 2017), with delayed larvae development (Wei et al., 2018). The effects of other pollutants, such as copper, on egg hormone transfer have not been addressed, yet the effects of pollutants on plasma levels (see above) suggest that such effects may be likely. However, no systematic investigation into the sources of variation in egg THs have been conducted in fish (Ruuskanen and Hsu, 2018).

We explored the causes of variation in maternal thyroid hormones in a common fish model, the three-spined stickleback (*Gasterosteus aculeatus*). Given the extremely scarce literature, we took an exploratory approach and studied environmental and potential genetic variation in maternal TH transfer using correlational and experimental approaches. This species was selected as it is an abundant and wide-spread species across the Northern Hemisphere, and an important species in biomonitoring and ecotoxicology as well as in behavioural and ecological studies (e.g. Scholz and Mayer, 2008). First, we sampled fish in six populations in the Baltic Sea (Fig 1), of which two were from discharge water areas of nuclear power plants, and four reference populations. Areas used for nuclear power discharge waters function as a natural experiment for long-term temperature increases, mimicking the effects of global warming, as discharged water is about 10-12 °C warmer than the intake water, and the discharge has continued for decades (Keskitalo and Ilus, 1987). This higher water temperature at the nuclear power plants could be a potential selecting agent on thermal physiology, including TH levels. We measured intraovarian THs of the fish from these six populations after an acclimation period in the laboratory. If there has been selection for altered TH metabolism and TH transfer in wild populations in these warmer water areas, we expect to see altered ovarian TH levels in the populations at the vicinity of the nuclear power plants compared to reference areas. Given that stickleback populations across the Baltic Sea are genetically different from each other, and the populations differ in response to thermal habitats and salinity they encounter (Guo et al., 2015), we may also expect overall differences in THs among the six populations. Furthermore, we tested the effects of temperature also experimentally by exposing the fish from each population to a mild temperature treatment (10°C increase), mimicking a heat wave. We predict that intraovarian T3 and T4 should be higher in the heat stress treatment compared to controls, due to increasing metabolic rates.

**Fig 1.**
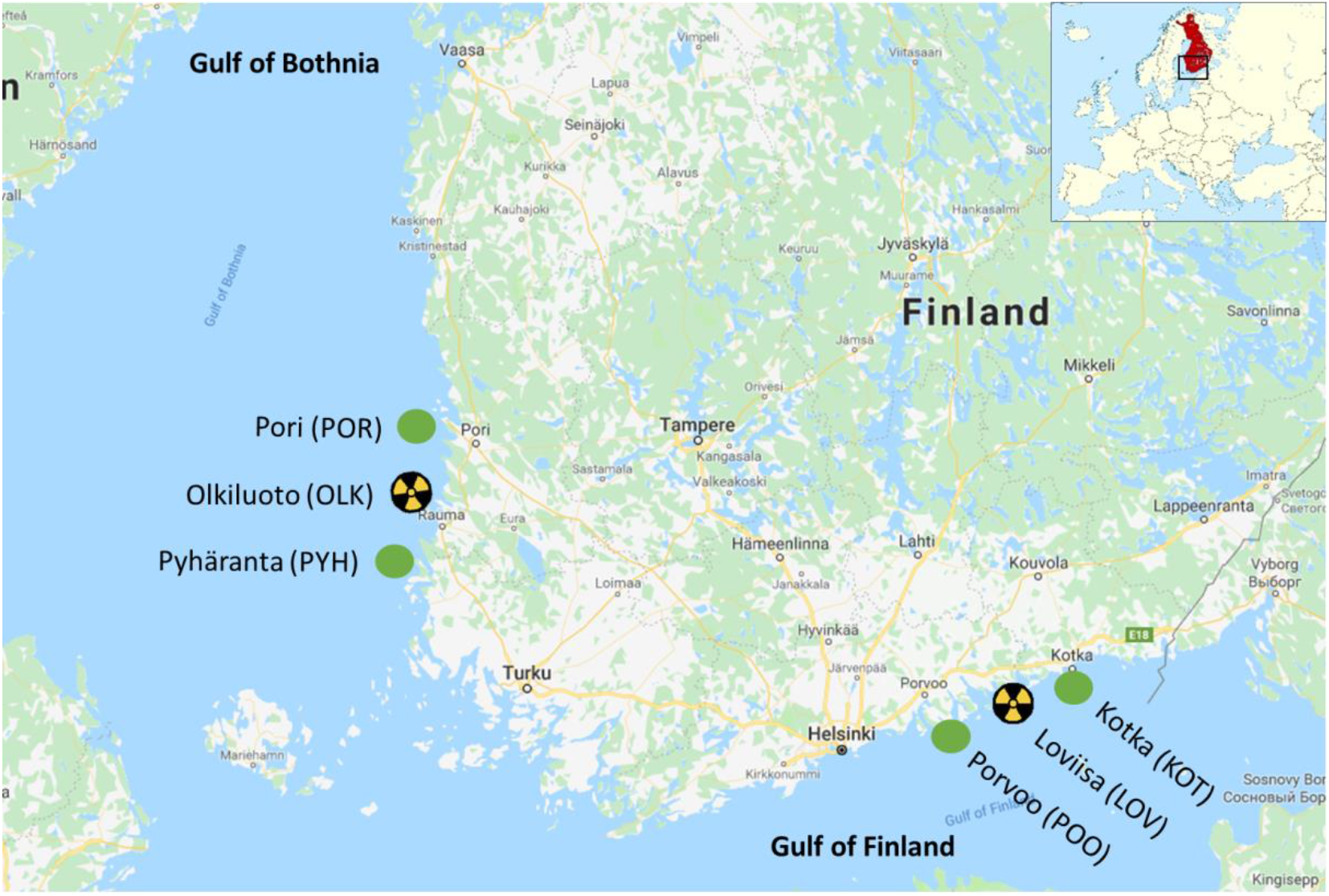
Sampling locations across the Baltic Sea. Yellow-and-black symbols refer to sites with nuclear power plant discharge water and green symbols to reference populations.

Second, contamination in the Baltic Sea is quite high with metals such as copper (but also zinc, lead, cadmium and mercury) reported in sediments, seawater, and e.g. liver tissues of herring and cod, and they influence e.g. physiology in mussels (Lehtonen et al., 2019; Perttila et al., 1982). Given that copper pollution can disrupt thyroid hormone levels in adult fish, we experimentally tested if maternal metal exposure to an environmentally relevant dose of copper can influence intraovarian hormone levels, i.e. allocation of hormones to eggs.

## Materials and methods

### Study areas, catching and maintenance

The experiments were conducted with wild, adult female three-spined sticklebacks caught from a wide geographical range, six different locations across the Baltic Sea (see Fig 1). Three of the locations were in Gulf of Finland and three in Gulf of Bothnia. In both Gulfs one of the locations was in the cooling water discharge area of a nuclear power plant, where discharged water is about 10-12 °C warmer than the surface water and the discharge has continued for decades (Keskitalo and Ilus, 1987). In Gulf of Finland the areas were Loviisa (LOV, Nuclear Power Plant area, N6021.928; E2622.228), Kotka (KOT, reference site, N6026.103; E2652.181) and Porvoo (POO, reference site, N6015.090; E2546.134). In Gulf of Bothnia the areas were Olkiluoto (OLK, Nuclear Power Plant area, N61.2360278; E21.4347222), Pyhäranta (PYH, reference site, N6057.149; E2125.986) and Pori (POR, reference site, N6130.140; E2135.675). Temperatures, salinity and pH of the locations during catching in May 2018 are presented in Supplementary Table 1. Temperature loggers (three per area, HOBO Water Temp Pro v2 Logger, U22, Onset Computer Corporation, Bourne, MA, USA) were situated directly at the catching locations at the sea bottom (depth 1.5-2 m) and water temperature was recorded 16/5/2018 – 31/8/2018 from all the locations four times per day. The average daily temperatures ± SD are presented in Supplementary Table 1.

The adult fish were caught with beach seine net and transferred to University of Turku for rearing. No mortalities were observed during the transfer and the acclimation period in laboratory facilities. The fish (N=100 per population, mixed sexes, some of the fish were used also in other experiments) were let to acclimate into 180 L tanks at +16°C for two weeks (each population in its own tank). Water salinity was 4 ppt (filtered water with 76% NaCl; 20% MgSO_4_; 3.5% CaCl_2_; 0.5% KHCO_3_), pH=8 and oxygen saturation over 80%. Photoperiod was set to 17L:7D. Fish were fed with frozen bloodworms (Delang & Ekman AB/ Akvarieteknik, Sweden) five times per week. One third of the water was changed once a week. Upon arrival the fish were treated against nematodes using Nematol (Sera GmbH, Heinberg, Germany) according to the instructions of the manufacturer. In order to reduce any tank effects the fish were tagged intraperitoneally with 1.35×7 mm RFID subcutaneous microchips (Loligo^®^ Systems, Viborg, Denmark) under anesthesia (100 ppm MS-222 in 4 ppt brackish water buffered with 6 ppm HCO_3_) after two weeks’ acclimation period. The populations were mixed into nine tanks (with density of 2L/fish) and were let to recover for three weeks before further testing.

### Experimental treatments

After the recovery period, the fish were exposed to three different environmental conditions in their rearing tanks: control (CTRL7D), sublethal level of copper (Cu 7D) or heat stress (HS 7D) for one week. Equal numbers of fish from each sampling location were distributed to all three treatments. Fish were sampled before and after the exposures (see below). The sub-lethal copper exposure (CU 7D) was conducted for a total number of 127 fish in three tanks (20 female fish from one tank were sampled for the current study). A total of 120 fish in three replicate control tanks (CTRL 7D) were treated similarly but no copper was added to tanks (25 female fish from one tank were sampled for the current study). For exposure, Cu^2+^ was added manually as copper (II) sulphate pentahydrate solution (nominal: 100μg/L of CuSO_4_·5H_2_O, Merck, Darmstadt, Germany) to the experimental tanks. This concentration of copper represents environmentally relevant concentrations encountered in polluted waters (Sanchez et al., 2005). Water samples were taken both from exposure and control tanks (i) 2 hrs and (ii) one week after the release of copper in order to measure the copper concentration during one week of exposure (no water changes were done during the exposure). Fifty millilitres of water was sampled in polypropylene Falcon tubes from tanks. In order to keep water samples fresh prior to analyses, 1 ml of concentrated HNO_3_/100ml was added to the samples and samples were kept at +4°C before analyses at SYNLAB Analytics & Services Finland Oy (Karkkila, Finland). The copper levels were measured by using inductively coupled plasma mass spectrometry (ICP-MS) (ThermoFisher Scientific, MA, USA). 5 fish died during the one week of exposure.

A third group of fish (N=125 in total in three tanks, 20 female fish from one tank were sampled for the current study) was exposed to heat stress (HS 7D) for a week during the same time as the other fish were exposed to copper/control treatment. For simulating an environmental heat wave the water temperature in the experimental tanks (16°C) was gradually increased by 1°C every 30 minutes until reaching 26°C and kept at this temperature for a week. The warming experimental set-up was done with chiller-heater (Julabo, Model: F32; AC: 230V 50/60 Hz 12A, Julabo GmbH, Seelback, Germany) connected to stainless steel coils. Six fish died during the exposure to heat stress.

### Sampling procedure

Before exposing fish to heat stress or copper, 17 female fish were sampled from a control tank (hereafter CRTL). Fish were sacrificed with cranial percussion. Tag code, weight (g), length (cm) were recorded from each fish. For the current study the ovaries were collected and flash frozen in liquid nitrogen. Samples were stored at −80°C for further molecular analyses. Similar sampling of the fish and ovaries was conducted also seven days after exposure for control (CTRL7D), copper exposure (Cu7D) and heat stress treatment (HS7D) for a total number of 65 female fish.

### Thyroid hormone analyses

The whole content of thawed ovaries (i.e. egg follicles and associated ovarian fluids) was gently squeezed out of the ovaries directly into a microcentrifuge tube and the sample was weighed (~0.001g). Samples were analysed for T3 and T4 at the University of Turku. LCMS/MS was conducted at the facilities of Turku Center for Biotechnology. T4 and T3 were extracted from yolk following previously published methods (de Escobar et al. 1985, Ruuskanen et al. 2018). In short, samples were homogenized in methanol in tissue lyser (Quiagen, Retsch GmbH, Haan, Germany). As an internal recovery tracer, a known amount of ^13^C_12_-T4 (Larodan, Sweden) was added to each sample. This allowed us to control for the variation in recovery (i.e. extraction efficiency) for each sample. Next, 400 μl of chloroform was added to sample. After centrifugation (15 min, 1900 g, +4°C), the supernatant was collected and the pellet was re-extracted in a mixture of chloroform and methanol (2:1). Back-extraction into an aqueous phase (0.05% CaCl_2_) was followed by a re-extraction with a mixture of chloroform:methanol: 0.05% CaCl_2_ (3:49:48) and this phase was further purified in-house on Bio-Rad AG 1-X2 (USA) resin columns. The iodothyronines were eluted with 70% acetic acid, and evaporated under N_2_.

Blanks (plain reagents without any sample) were analysed in each extraction batch to detect any contamination. Samples from different populations and treatments were equally distributed across four extraction batches. There was no difference among the batches in hormone concentrations (F < 1.0, p > 0.38). T3 and T4 were quantified using a nanoflow liquid chromatography-mass spectrometry (nano-LC-MS/MS) method, developed and validated in Ruuskanen et al. (2018). Briefly, before the analysis, the dry samples were diluted in ammonium (NH_3_). Internal standards ^13^C_6_-T_3_ and ^13^C_6_-T_4_ (Sigma-Adrich, St.Louis, USA) were added to each sample to identify and quantify the THs. A triple quadrupole mass spectrometer (TSQ Vantage, Thermo Scientific, San Jose, CA) was used to analyse the samples. For the chromatographic separation of hormones, a nanoflow HPLC system Easy-nLC (Thermo Scientific) was applied. On-column quantification limits were 10.6 amol for T4 and 17.9 amol for T3. MS data was acquired automatically using Thermo Xcalibur software (Thermo Fisher Scientific) and analysed using Skyline (MacLean et al. 2010). For the analyses, peak area ratios of sample to internal standard were calculated. TH concentrations are expressed as pg/mg fresh mass. Few samples failed in the extraction, see samples sizes in Fig 3.

### Statistitcal analysis

All statistical analyses were conducted with SAS Enterprise Guide version 7.1. T4 concentration (pg/mg) was log-transformed to reach normality. We first analysed differences in intraovarian T3 and T4 among long-term heat exposed (i.e. populations in the vicinity of the nuclear power plants that are exposed to warm discharge waters; pooled LOV and OLK, N = 13 individuals) and reference populations (pooled POO, POR, KOT, PYH, N = 24 individuals) using a linear model, using fish mass as a covariate. Data from individuals from control treatments (CRTL and CRTL7D) were only included in this analysis. We then analysed differences in intraovarian T3 and T4 concentration among all populations, again using data only from the control treatments. From POO, we only had 1 individual for this analysis, and thus this population was excluded from the analysis. Population and fish mass were included as fixed effects.

Differences in ovarian T3 and T4 concentration between treatments (CRTL 7D, Cu 7D, HS 7D) were analysed using linear mixed models where treatment and fish body mass were included as fixed effects and population as a random intercept to account for non-independence of fish from the same population. Finally, we also repeated the above models for fish size (mm). Post-hoc tests were further used to test pair-wise differences among treatments. Models were reduced by removing non-significant factors (α = 0.05). Normality and heteroscedasticity of the residuals were visually inspected. Degrees of freedom were estimated with Satterthwaite estimation method. Means and standard errors (SE) are shown in the text and in the figures.

### Results

When using data from control treatments only (pooled CRTL and CRTL 7D), we found no differences in intraovarian T3 or T4 concentration in sites exposed to warm discharge waters (pooled LOV and OLK) compared to reference sites (pooled four reference sites, T3: F_1,36_ = 0.88, p = 0.18, T4: F_1,36_ = 0.25, p = 0.62; Mean (pg/mg) ±SE: T3 exposed populations: 1.60±0.19, T3 reference populations: 2.21±0.24; T4 exposed populations: 0.57±0.08; T4 reference populations 0.53±0.04). When further comparing all populations, we found no strong statistical evidence for differences among populations in intraovarian T3 or T4 concentration (T3: F_4,30_ = 0.50, p = 0.73, T4: F_4,30_ = 2.38, p = 0.08; Fig 2a, b). Body mass correlated negatively with intraovarian T3 concentration (estimate±SE: −0.842±0.363, F_1,30_ = 5.37, p = 0.027), but not with T4 concentration (F_1,30_ = 0.0, p = 0.96). Fish size did not differ among exposed and reference populations (F_1,35_ = 1.89, p = 0.18), while there were small differences among populations (F_4,32_ = 6.92, p = 0.004): fish from KOT were significantly smaller than those from OLK, PYH and POR (see Suppl Table 2).

**Fig. 2.**
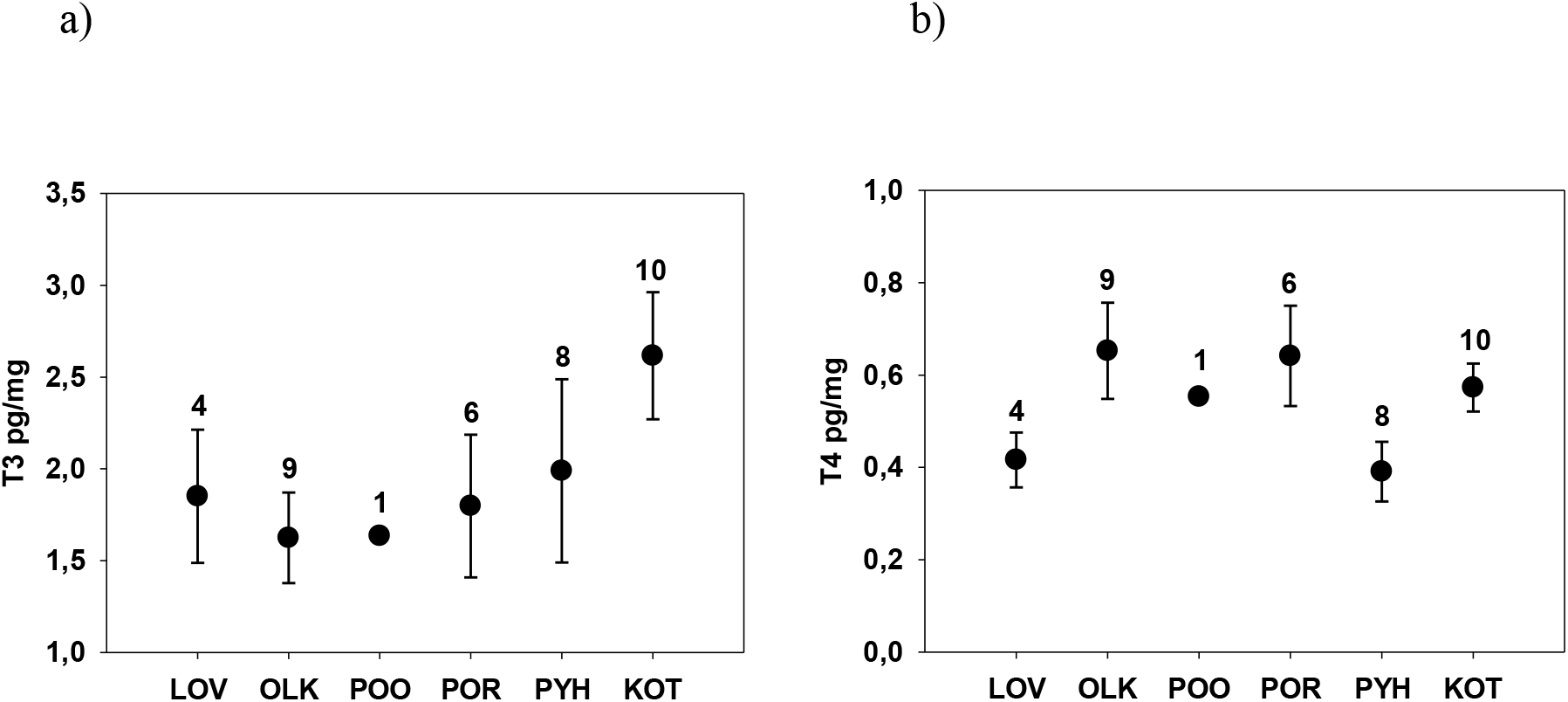
Intraovarian T3 (a) and T4 (b) concentrations (pg/mg, mean±SEs) in three-spined sticklebacks from six populations across the Baltic Sea coast. Numbers above the bars refer to sample sizes. POO was excluded from the statistical analysis due to low sample size.

In the copper exposure treatment, the measured concentrations of copper were 91, 101, 91 and 22 μg/L after 2 hrs and 35, 48, 37 and 18 μg/L after one week in the three exposure and one control tank, respectively. The exposure concentration was relatively high but within environmentally relevant concentration range and has not been shown to cause mortality in sticklebacks in previous studies (Sanchez et al., 2005). According to chemical water analyses all the fish in current study were exposed to low concentration of the copper since they were brought to laboratory facilities due to the technical purity of salts for producing brackish water. After experimental exposure to copper for 7 days, fish from copper exposure group had higher intraovarian T3 and T4 concentration compared to fish from control treatment (T3: overall test F_2,53.1_ = 3.14, p = 0.05, post-hoc Control vs Cu t_52.3_ = −2.30, p = 0.02; T4: F_2,53_ = 2.62, p = 0.08, Control vs Cu t_53.8_ = −2.26, p = 0.027, Fig 3a, b). T3 or T4 concentrations of fish from the heat treatment did not differ statistically from control treatment (post-hoc p-values >0.45), while ovarian T3 concentration was lower in heat treatment compared to copper exposed fish (t_52_ = 2.52, p = 0.045, Fig 2a). Fish size did not differ among the treatments (F_2,51_ = 0.64, p = 0.53).

**Fig. 3.**
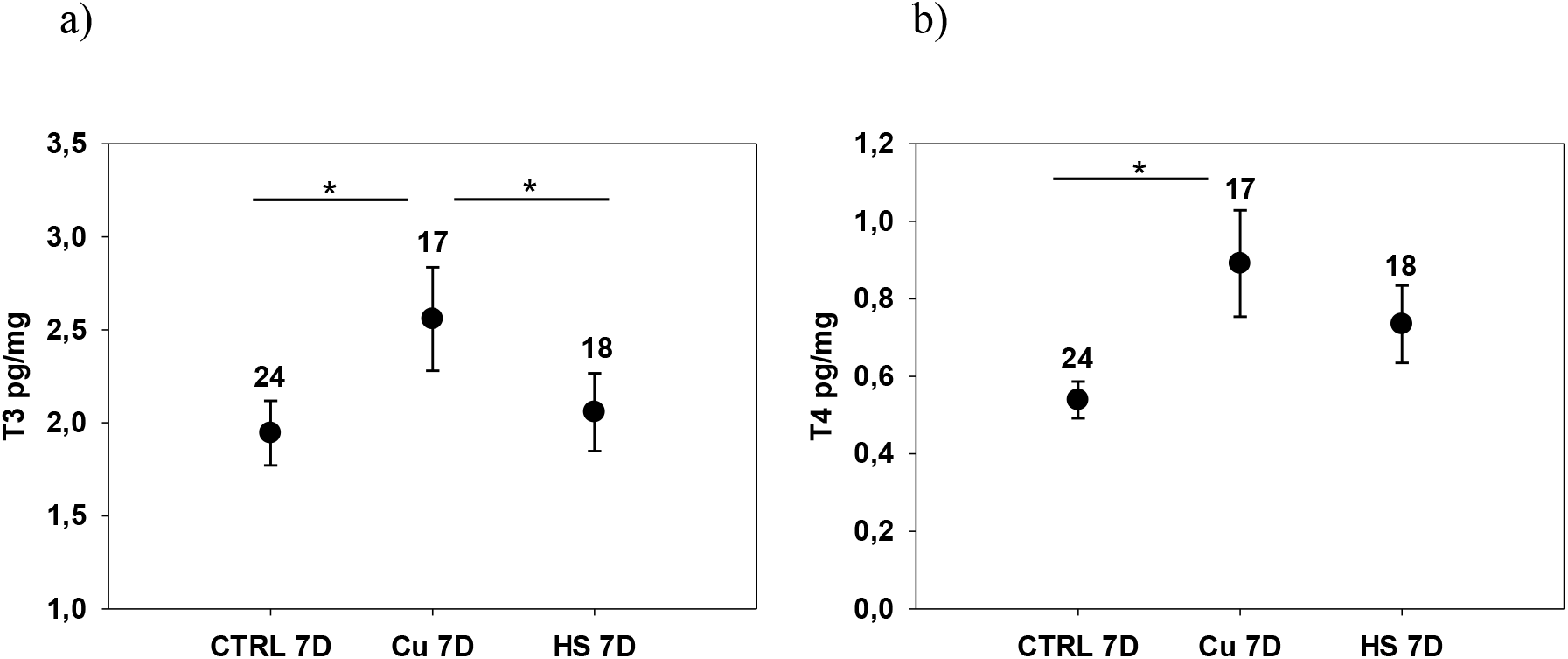
Intraovarian T3 (a) and T4 (b) concentrations (pg/mg, mean±SEs) in three-spined sticklebacks experimentally exposed to copper (100μg/L, CU 7D), warm temperature (10°C increase, HS 7D) or respective controls (CRTL 7D) for 7 days. Numbers above the bars refer to sample sizes. Stars represent significant differences between treatments at p < 0.05. For the statistical analyses the populations were pooled.

## Discussion

In contrast to our predictions, we found no evidence that intraovarian TH levels of individuals originating from populations close to a long-term heat source (nuclear power plant discharge waters), differ from individuals originating from reference populations. Furthermore, the six populations sampled across a wide geographical range over the Baltic Sea showed similar intraovarian TH levels after acclimation in captivity. These results suggest that populations do not show strong genetic (or permanent developmentally induced) variation in intraovarian T4 or T3 levels, although at least some Baltic Sea stickleback populations are known to be genetically differentiated (Guo et al., 2015). However, we cannot rule out that environmental sources of variation (temperature, salinity, food availability) could influence intraovarian THs in the wild. Among-population variation in egg THs in field populations was indeed reported by McComb et al. (2005) in bonnethead sharks. Nevertheless, given that intraovarian THs were not influenced by short-term experimental exposure to heat, we may conclude that temperature – using the tested experimental duration and scope – may not influence transfer of THs to eggs and offspring. The sample size was, however, rather low in the among-population comparison, and the differences in environmental conditions (see Suppl Table 1) among the populations were rather small at the time of the sampling, thus the results should be interpreted with caution. Interestingly, stickleback adaptive divergence from marine to stream environment has been found to involve thyroid hormone signalling (Kitano and Lema, 2013; Kitano et al., 2010), thus it would be interesting to further compare the egg THs from freshwater and marine populations to understand the role of transgenerational plasticity in the adaptation process.

We found that individuals exposed to environmentally relevant concentrations of copper for seven days showed higher intraovarian T3 and T4 levels than controls. While associations between copper and maternal THs have not been studied to date in fish, previous studies on the associations between copper as an EDC and THs in plasma show complex patterns depending on the duration of the exposure and species, as reported in Eyckmanns et al. (2010): In common carp (*Cyprinus carpio*), T3 levels were *elevated* only after long-term exposure (1 month), while in gibel carp (*Carassius gibelio*) there was a *decrease* in T3 from 24 h to 1 month of exposure. Both species showed *increases* in T4 over short-and long-term exposure. In rainbow trout, T4 levels were *elevated* very fast after copper exposure and remained elevated for 12h, whereas there was *no influence* on T3. Copper exposure also *increased* plasma T4 in the common carp in another experiment (Hoseini et al., 2016). Finally, copper exposure significantly *decreased* T3 but not T4 in European eels (*Anguilla Anguilla*) (Oliveira et al., 2008), suggesting changes in deionization from T4 to T3 in tissues. These inconsistencies may be explained by experimental conditions and dosages, exposure time and species-specific responses.

The association between plasma and intraovarian/egg hormone levels in fish has not been fully elucidated, but if we assume that there is some correlation between intraovarian and circulating TH levels (Ayson and Lam, 1993; Brown et al., 1988; Brown et al., 2014; Kang and Chang, 2004; Raine and Leatherland, 2003), our results are in parallel with those of common carp (see above). Interestingly, our hormone measurements of the two forms (T3 higher than T4) were contrasting compared to whole body measurements in the same species (T3 lower than T4, Gardell et al., 2017), and egg measurements on other species (T3 lower than T4, Chen et al., 2017). We can speculate that this may be due to differential deposition of the two forms, or deiodinase function, converting T4 to T3 in the ovaries in these species and sample type. The increased T4 and T3 levels in response to copper exposure reported in this study suggest that T4 biosynthesis or degradation was altered, but also that conversion of T4 to T3 was potentially increased. Increased TH levels could be explained by increased metabolic rates and energy expenditure in response to copper exposure, along with increased oxidative stress (De Boeck et al., 2006; De Boeck et al., 1997; Sanchez et al., 2005). All in all, this evidence suggests that copper exposure changes plasma and associated ovarian TH levels. The next step would be to evaluate whether such changes lead to changes in on offspring development, such as metabolism and thermotolerance.

In contrast to our predications, we did not find any differences in TH concentrations between experimental heat treatment and control. In a previous study it has been found that three weeks acclimation to high and low temperature changed the muscle TH profile of zebrafish, warm acclimated fish having higher concentrations of T3 and T2 than cold acclimated ones (Little et al., 2013). However, in their study the warm-acclimated fish were not sensitive to changes in hormone levels suggesting that with temperature exposure also e.g. TH transporters and receptors need to be evaluated. This could be one reason why we did not see any change in egg TH levels with heat stress even though it is well known that heat stress increases the metabolic rate of fish (e.g. Anttila et al., 2013; Fry, 1947).

We conclude that variation in intraovarian THs was not explained by (genetic) variation among populations nor short-term heat exposure. Given that parental temperature environment is known to alter offspring phenotype (e.g. Donelson et al., 2012; Donelson et al., 2018), further studies are needed to elucidate the molecular mechanism of such transgenerational effects. Both T4 and T3 levels in ovaries were altered in response to moderate copper exposure, and now the next step is to characterize potential functional consequences of altered THs on offspring phenotype, which would allow us to understand the scope for transgenerational endocrine disruption.

## Compliance with Ethical Standards

All procedures were conducted under licenses from the Animal Experiment Board of the Administrative Agency of South Finland (ESAVI/2867/2018). All applicable international, national, and/or institutional guidelines for the care and use of animals were followed. A pre-print version of this manuscript has been submitted to BioRxiv (Ruuskanen et al. 2019) and will be replaced with the final version upon acceptance.

## Acknowledgements

We thank Jenni Saukkonen and Tytti Uurasmaa for field and experimental support.

---

**Supplementary Table S1.** The environmental variables (mean±SD) of catching locations across the Baltic Sea.

| Location | Temperature<br>during catching | Salinity | pH | Average ± SD temperature<br>16/5/–31/8/2018 |
| --- | --- | --- | --- | --- |
| LOV | 19.9 ± 1.6 | 3.2 ± 0.2 | 7.6 | 19.6 ± 4.4 |
| OLK | 20.1 ± 0.1 | 5.3 ± 0.0 | 9.0 | 19.9 ± 3.6 |
| POO | 18.8 ± 0.5 | 4.3 ± 0.0 | 9.0 | 19.1 ± 4.2 |
| POR | 18.9 ± 0.8 | 4.9 ± 0.0 | 9.0 | 19.7 ± 3.4 |
| PYH | 17.6 ± 1.1 | 4.2 ± 0.4 | 9.0 | 19.5 ± 2.5 |
| KOT | 18.1 ± 0.7 | 1.7 ± 0.0 | 9.0 | 17.3 ± 4.9 |

**Supplementary Table S2.**
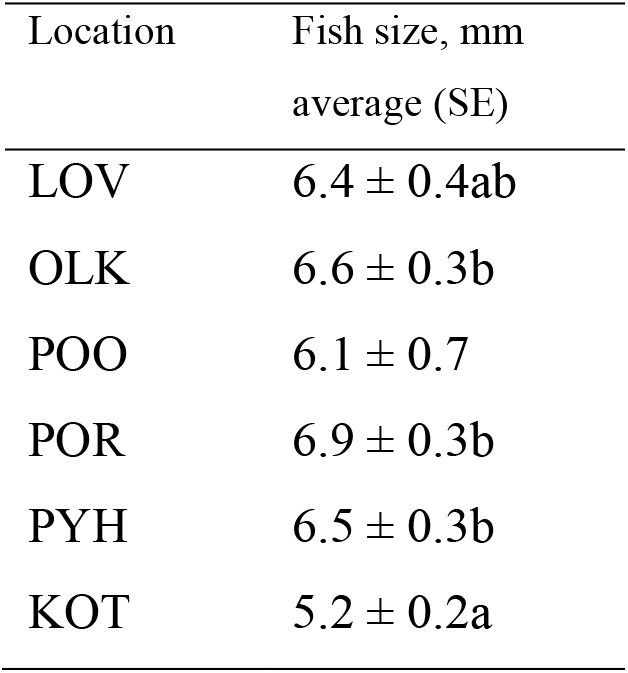
The average lengths (mm, SE) of fish captured from six different locations (populations) across the Baltic Sea. Populations with different letters are statistically significantly different from each other (p <0.05, Tukey post-hoc test). POO was not included in the statistical analyses due to small sample size.

